# An Algorithm to Quantify Inducible Protein Condensates In Eukaryotic Cells

**DOI:** 10.1101/2021.08.26.457826

**Authors:** Jeremy C. Hunn, Katherine M. Hutchinson, Joshua B. Kelley, Daniel Reines

**Author notes:** These author contributed equally to this work (Jeremy C. Hunn, Katherine M. Hutchinson).

## Abstract

Reorganization of cellular proteins into subcellular compartments, such as the rearrangement of RNA-binding proteins into cytoplasmic stress granules and P-bodies, is a well-recognized, widely studied physiological process currently under intense investigation. Using the assembly of a novel, inducible, nuclear granule formed from the yeast RNA-binding transcription termination factors Nab3 and Nrd1, we present a freely-accessible, high-throughput and unbiased algorithm written in MATLAB that detects and measures protein distribution, partitioning, and sequestration into subcellular compartments captured by fluorescence microscopy; an invaluable advancement to current image analysis methods which utilize experiment-specific custom scripts or subjective manual counting. Employing our algorithm, we quantified thousands of cells, ensuring rigorous examination of Nab3 granule formation across strains with reproducible statistical analyses. We document strain differences in Nab3 granule formation and an associated growth defect. Additionally, we applied our algorithm to immunofluorescent images of the inducible polymerization into filaments of an enzyme in human cells, demonstrating the algorithm’s versatility and adaptability.

**SUMMARY STATEMENT:** We describe a computational tool that enables the quantification of protein condensation during assembly of a subnuclear compartment. The algorithm scores assembly of fluorescently tagged proteins in yeast or human cells.

## INTRODUCTION

The reorganization of proteins in response to physiological stimuli to form subcellular compartments, such as stress granules and P-bodies, has provided novel insight into cellular structure and the mechanisms by which cells compartmentalize their biochemical and molecular events. RNA-binding proteins, such as Pub1 found in yeast stress granules (Lin et al., 2015), are frequently examined for their protein reorganization as this class of protein often harbors low complexity domains (LCD), making them capable of self-assembly (Harrison and Shorter, 2017; Weber and Brangwynne, 2012). LCDs typically lack stable secondary structural elements due to their intrinsically disordered nature; however, the domain can become structured and highly organized by assembling via a phase separation mechanism. Phase separation and protein rearrangement are studied by a variety of methods, fluorescence microscopy being one well established modality (Alberti et al., 2019).

The advent of fluorescent protein tags in combination with improvements in microscope technology has enabled researchers to study protein localization and reorganization in great detail, such as the inducible rearrangement of proteins involved in transcription and RNA metabolism in response to physiological stimuli. These technological advancements create a need for a high throughput method to quantify the rearrangement of protein captured by fluorescent microscopy. Current methods to quantify and analyze images can be tedious and time consuming, often requiring researchers to manually count blinded images or deploy custom scripts written specifically to work with their data on various software platforms that are not always publicly accessible. As the field of fluorescent microscopy continues to advance and evolve, the demand for a readily accessible, adaptable and versatile algorithm persists.

To address this demand, we created an algorithm that uses single-cell fluorescence histogram analysis to quantify protein rearrangement captured by fluorescence microscopy. We utilized the essential *S. cerevisiae* RNA-binding transcription factors, Nab3 and Nrd1, as a paradigm to demonstrate the capabilities of a granule-calling algorithm. Nab3 and Nrd1 are RNA-binding proteins that contain glutamine-rich low complexity domains (Alberti et al., 2009; Loya et al., 2013b; Loya et al., 2018) as well as an RNA recognition motif, which mediates RNA-binding. Through binding to each other, RNA, and RNA polymerase II-binding, they mediate the termination of RNA polymerase II transcription for many yeast transcripts, particularly small non-coding RNAs such as snRNAs and snoRNAs, as well as select mRNAs (Arndt and Reines, 2015; Creamer et al., 2011; Jamonnak et al., 2011; Jenks et al., 2008; Kopcewicz et al., 2007; Loya et al., 2013a). Nab3 and Nrd1 are excellent candidates for studying protein rearrangement as these proteins reorganize into tight puncta in nuclei when cells are deprived of glucose as a carbon source (Darby et al., 2012). Assembly requires the Nab3 LCD and is rapidly and fully reversible when cells are fed metabolizable sugar (Loya et al., 2018). This punctate structure is novel as Nab3 does not colocalize with the stress granule marker Pub1, the P-body marker Dcp2, or the nucleolar marker Nop1, (Darby et al., 2012) making it a new class of subcellular structure that contains transcription termination factors but whose function is poorly understood.

Through the use of our algorithm, we have quantified and characterized the extent of Nab3 granule formation in three strains of *S. cerevisiae*. For the first time, we demonstrate that Nab3 granule formation in response to glucose deprivation varies across *S. cerevisiae* strains. Furthermore, we document a growth defect that correlates with the granule forming phenotype, raising the possibility that Nab3 granule formation is related to cell signaling and proliferation. While the method enabled us to measure a number of parameters of nuclear granule formation in live cells, we also show that the method is generally applicable since it can also be used to score protein assemblies detected in immunofluorescent images of fixed mammalian cells.

## RESULTS

### Design of an Algorithm for Granule Detection

Glucose starvation in yeast is a well-established stimulus used to provoke the formation of stress granules and P-bodies which are cytoplasmic compartments that harbor RNA and RNA-binding proteins (Buchan et al., 2010). Glucose restriction results in a change in steady state mRNA composition (Zaman et al., 2008) and a physical redistribution of the transcription termination factors Nab3 and Nrd1 from a pan-nuclear distribution to tight nuclear granules (Darby et al., 2012). This dynamic re-localization is accompanied by a change in the RNA-binding profiles of these termination factors (Bresson et al., 2017; Creamer et al., 2011; Jamonnak et al., 2011). The initial finding of this redistribution and a more recent confirmation using confocal microscopy (Loya et al., 2018), employed manual counting to quantify the fraction of cells bearing the Nab3-Nrd1 granule. Our previous work also used a computational approach to detect granule formation over time in single cell time-lapse microscopy. However, the previous algorithm relied upon kinetic changes and was not applicable to other types of data sets. During our investigation of this process, we noted varying levels of Nab3 granule formation following glucose starvation in large numbers of cells at a single time point across distinct yeast strains (Fig. 1). Thus, we sought to develop an automated and unbiased method to quantify Nab3-containing granules in thousands of cells from multiple biological replicates that was not limited to time-lapse microscopy.

**Figure 1.**
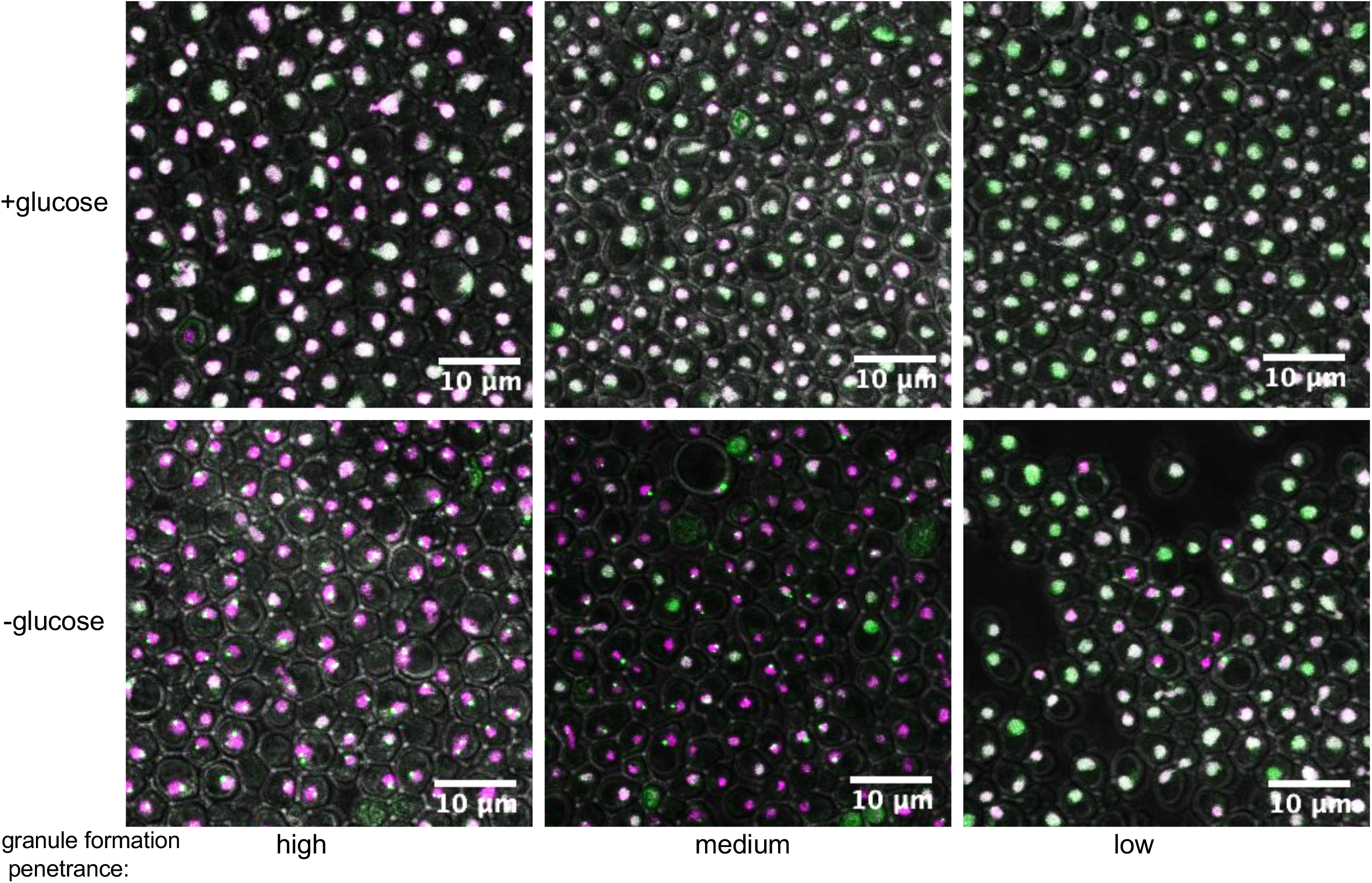
Fluorescence microscopy of GFP-Nab3 (green) granule formation in three yeast strains. Three different strains of yeast cells were grown to mid-log phase, washed free of growth media, and resuspended in fresh media with or without glucose as described in Materials and Methods. Nuclei were labeled with histone H2B tagged with mCherry. Nab3 was tagged with GFP. (r.f.u.= relative fluorescent units).

Three yeast strains were each engineered to contain two fluorescent fusion proteins, a red fluorescent protein fused to histone H2B, and Nab3 tagged with green fluorescent protein (GFP) (Table 1). The former serves to mark the nucleus and the latter is the protein of interest. Both fusion genes reside at their natural chromosomal loci with expression driven by their native promoters. One strain is a standard laboratory strain of *S. cerevisiae* (BY4741) that is a derivative of S288C (Brachmann et al., 1998). Another strain is the laboratory strain W303 (Horizon Discovery, LLC). The third strain is a variant of W303 engineered to contain NRD1 on a plasmid covering a chromosomal deletion of the gene. Cells in exponential growth were washed and incubated for 2 hr at 30°C in media lacking glucose, applied to an agarose pad, and examined using a Leica SP8 multiphoton microscope. The microscope-native .lif file format was converted to a MATLAB readable file (in this case, a .png file) and analyzed by a custom MATLAB algorithm (available through GitHub at https://github.com/Kelley-Lab-Computational-Biology/Granule) that scores the redistribution of fluorescence intensity in cells before and after granule formation.

**TABLE 1.**

| Strain | Genotype | Reference |
| --- | --- | --- |
| BY4741 | *MATa his3Δ1 leu2Δ0 met15Δ0 ura3Δ0* | Research Genetics |
| DY1620 | *MATα his3-11,15 leu2-3,112 trp1-1 ura3-1 NRD1::natMX4* [pAC3314-Nrd1] | A. Corbett (Emory U.) |
| DY4543 | *MATa his3Δ1 leu2Δ0 met15Δ0 ura3Δ0 GFP-NAB3* | Loya & Reines, 2018 |
| DY4580 | *MATa his3Δ1 leu2Δ0 met15Δ0 ura3Δ0 GFP-NAB3 HTB2-mCherry::HIS3* | This Study |
| DY4724 | *MATα his3-11,15 leu2-3,112 trp1-1 ura3-1 NRD1::natMX4* [pAC3314-Nrd1] [pRS315-HA-Nrd1] | This Study |
| DY4736 | *MATα his3-11,15 leu2-3,112 trp1-1 ura3-1 NRD1::natMX4* [pRS315-HA-Nrd1] | This Study |
| DY4746 | *MATα his3-11,15 leu2-3,112 trp1-1 ura3-1 NRD1::natMX4 GFP-NAB3* [pRS315-HA-Nrd1] | This Study |
| DY4756 | *MATα his3-11,15 leu2-3,112 trp1-1 ura3-1 NRD1::natMX4 GFP-NAB3 HTB2-yomCherry::kanMX* [pRS315-HA-Nrd1] | This Study |
| DY4771 | *MATα ade2-1 can1-100 his3-11,15 leu2-3,112 trp1-1 ura3-1 GFP-NAB3* | This Study |
| DY4772 | *MATα ade2-1 can1-100 his3-11,15 leu2-3,112 trp1-1 ura3-1 GFP-NAB3 HTB2-mCherry::HIS3* | This Study |
| DY4789 | *MATa/ MATα ADE2/ade2-1 CAN1/can1-100 his3Δ1/ his3-11,15 leu2Δ0/ leu2-3,112 met15Δ0/MET15 TRP1/trp1-1 ura3Δ0/ura3-1 GFP-NAB3/GFP-NAB3 HTB2-mCherry::HIS3/ HTB2-mCherry::HIS3* | This Study |
| YOL890 | *ura3-1 his3-11,15 leu2-3,112 lys2 htb2-mCherry::HIS3* | S. Wente (Vanderbilt U.) |
| YSC1059 | *MATα ade2-1 can1-100 his3-11,15 leu2-3,112 trp1-1 ura3-1* | Horizon Discovery LLC |

The workflow for the algorithm is shown in Fig. 2. First, nuclei are identified using both intensity and size criteria. We chose a fluorescence threshold that identified the vast majority of nuclei (extremely dim outliers were excluded). We also applied a size filter that excluded smaller objects that passed through the fluorescence filter but were not actually nuclei. We used a fluorescence value of 17 for the threshold, and a minimum size of 40, but these values are likely to be specific to our data set. The objects that met these criteria were considered nuclear masks. Within a mask, the fluorescent intensity of pixels with GFP-Nab3 were averaged after excluding the 10 highest value pixels. This exclusion was done to prevent skewing of the average value with the bright pixels that would be present in potential granules. The degree to which this is necessary will vary based on the total fraction of the protein that goes into a granule, and the number of pixels to exclude will depend on the size of the granules in the data set. The mean and standard deviation (σ) of pixels in the mask were calculated and pixels with intensities >3σ (“3σ” pixels) over the mean were identified. We have found that 3 times the standard deviation works well, but there may be some data sets or systems where a different range is desirable. This can be adjusted in the program. The size of clusters, in terms of the number of adjacent 3σ pixels, was scored for each nuclear mask.

**Figure 2.**
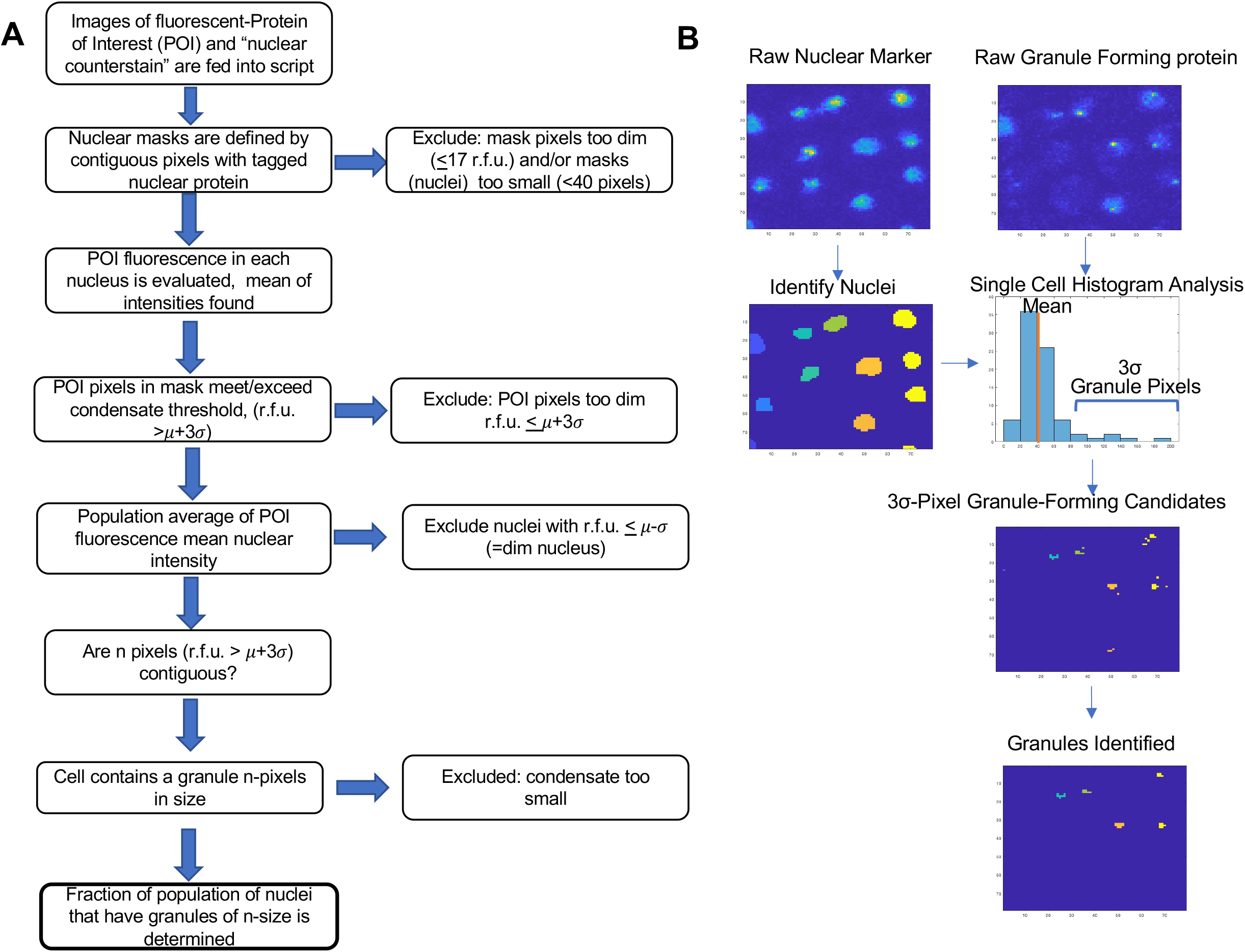
Workflow schematic. A. The steps employed to process digital micrographs are shown in flow chart form. The individual steps are described in the text. B. Images from the workflow for the MATLAB algorithm showing steps in image processing.

### Quantification of induced GFP-Nab3 rearrangement in yeast nuclei using digitized images

A primary consideration in the quantitation of these granules is the size of objects that can be considered a granule. Allowing any single pixel of sufficient intensity to be counted may identify noise as a granule. However, requiring an object to be particularly large in order to be scored as a granule may miss actual granules. Quantification that is overly sensitive to criteria risks not being robust across different conditions and samples. To address this issue, we examined granule formation across a range of size cutoffs. We first characterized the algorithm using a strain whose population displayed a very high level of cells with granules under starvation conditions (Fig. 1, “high”). To choose how large an object would be considered a granule, we plotted the percent of nuclei containing pixel clusters versus minimum granule size (Fig. 3A). In other words, the graph displays the fraction of cells defined as containing a granule for each minimum size of a potential granule. Glucose-fed cells do not contain granules and rarely, if ever, contain a cluster of 3σ pixels larger than three. Whereas a vast majority of glucose starved cells contained clusters of three or more of these very intense pixels. Almost half of the starved cells contained very large clusters of GPF-Nab3 fluorescence (≥8 pixels).

**Figure 3.**
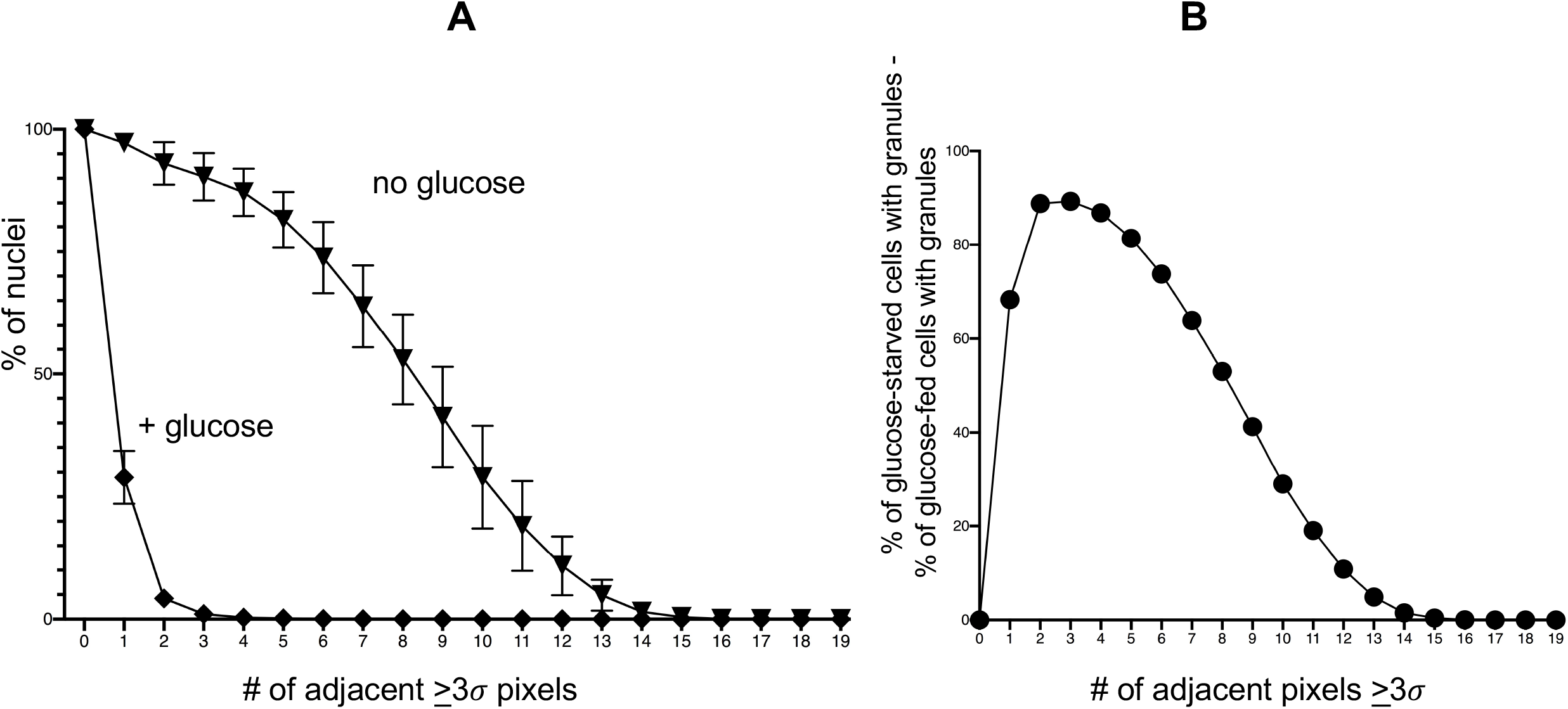
Measurement of the distribution of nuclear GFP-Nab3 in glucose-fed and glucose-starved conditions for a high-efficiency granule forming strain. A. *Clustering of 3σ pixels in yeast nuclei before and after glucose starvation*. Fluorescent images were analyzed by a MATLAB algorithm to score pixel intensity of GFP-Nab3 in a nuclear mask defined by H2B-mCherry. The number of adjacent pixels whose intensities were greater than or equal to three standard deviations (3σ) above the mean pixel intensity in that mask, were scored. The percent of cells (Y-axis) bearing a given number of 3σ pixel clusters (X-axis) was analyzed for six biological repeats (n=6; different cultures on different days) and the mean and standard deviations were calculated. The Y-axis value for zero adjacent ≥ 3σ pixels was set at 100%. B. *Difference between glucose-fed and glucose-starved cells in terms of their GFP-Nab3’s distribution at each 3σ pixel cluster size*. The “% of cells” values (Y-axis in part A) at each pixel cluster size (X-axis in part A) in glucose-fed cells from part A were subtracted from their Y-axis value at that pixel cluster size for the cognate glucose-starved cells. This difference was then plotted for each pixel cluster size.

As a result of this analysis, we chose to define a Nab3 granule as a cluster of ≥4 adjacent 3σ pixels. This was guided by the finding that the maximal difference between glucose-depleted and glucose-fed cultures, in terms of the percent of cells harboring clusters of 3σ pixels, was observed at ≥3 pixels (Fig. 3B).

### The algorithm enables a detailed analysis of fluorescent protein distribution

By eye, it appeared that almost all nuclei in this strain contained only a single granule. To quantify this, we plotted the number of cells that contained 1, 2, or 3 granules per nucleus (Fig. 4A). This analysis shows that nuclei with granules typically contain only a single granule and only rarely (5.8+/−5.3%) contain a second granule. Thus, the granule most often appears as a single subcellular compartment as opposed to multiple compartments within the nucleus.

**Figure 4.**
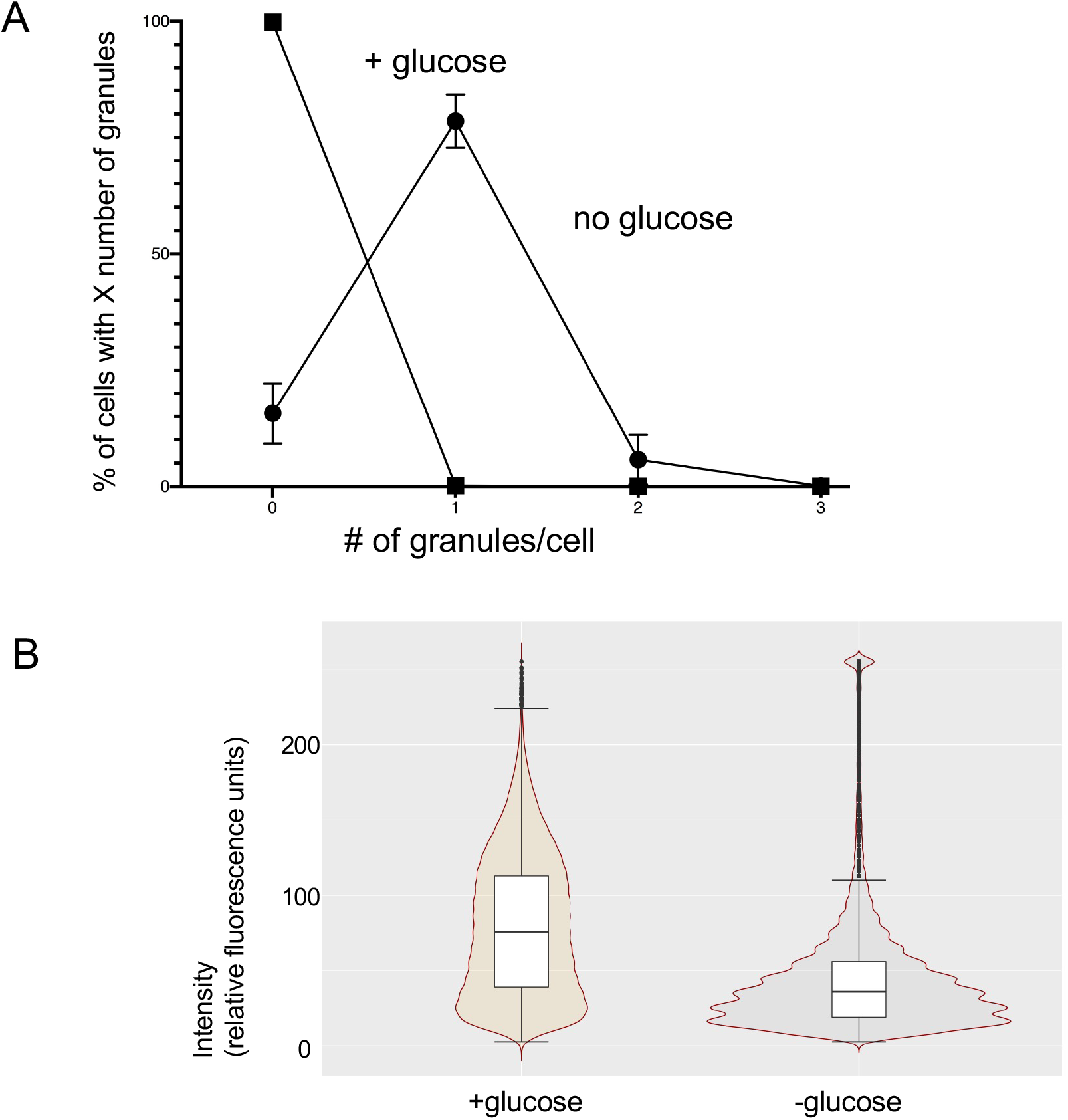
Quantification of granules per cell and distribution of pixel intensities for glucose-fed or glucose-restricted cells. A. *Granules per cell*. Yeast strain DY4756 was grown in the presence or absence of glucose. The number of granules of ≥4 pixels with ≥3σ intensities above the mean was calculated for each nucleus in cells from each of six biological replicates (n=6). Averages and standard deviations were plotted. B. *Distribution of pixel intensities for populations of fed or starved cells*. All values for GFP-Nab3 pixel intensities from a single field each of strain DY4756 grown in the presence or absence of glucose was collected. From these datasets, 100,000 pixels were randomly selected from each sample in MATLAB using the datasample function (without replacement). The datasets were then imported into RStudio and violin plots were created using the ggplot2 package, scaled by area.

To gain a better understanding of the distribution of GFP-Nab3 across a population, we plotted the fluorescence intensity of 100,000 randomly chosen nuclear pixels from a single field each of cells from glucose-replete and glucose-depleted cultures. As seen in the violin plot of Fig. 4B, the median fluorescence intensity was 76 relative fluorescence units for glucose-fed cells with a very broad spread of values across the 8-bit intensity spectrum (0-255). Glucose-depletion dropped the median substantially to 36 units where most nuclear pixels had very low fluorescent values, while a small number of pixels gained a very high fluorescence score as GFP-Nab3 collected in tight foci. The data captured by the algorithm emphasizes that GFP-Nab3 undergoes a dramatic rearrangement in which it condenses to a small nuclear volume during glucose starvation, at the expense of the broadly nuclear distribution seen under vegetative growth conditions.

We also plotted the pixel intensity distribution for specific glucose-fed and glucose-starved cells to emphasize the distribution of GFP-Nab3 in individual nuclei (Fig. 5). Six randomly chosen 100-pixel masks were analyzed and plotted for both glucose-fed and glucose-depleted cells. This analysis again shows that the more intense 3σ pixels (Fig. 5, red) draw fluorescence from the majority of pixels, driving them to a more compact collection of low intensity pixels. It also emphasizes that the even distribution of pan-nuclear GFP-Nab3 in vegetatively growing cells, rarely present as 3σ pixels, confirming the stringency of this cutoff. The quantification, on a per nucleus basis, of the fraction of GFP-Nab3 that migrates to a 3σ value upon starvation is also quantifiable for these representative nuclei, varying from 25-44%. When a similar analysis is carried out for all nuclei in an entire field, an average of 26% (+/−14% standard deviation, n=1121 cells) of GFP-Nab3 ends up in the granule after glucose restriction, while 1% (+/−2%, n=1264 cells) of the protein finds itself in 3σ pixels in glucose fed cells.

**Figure 5.**
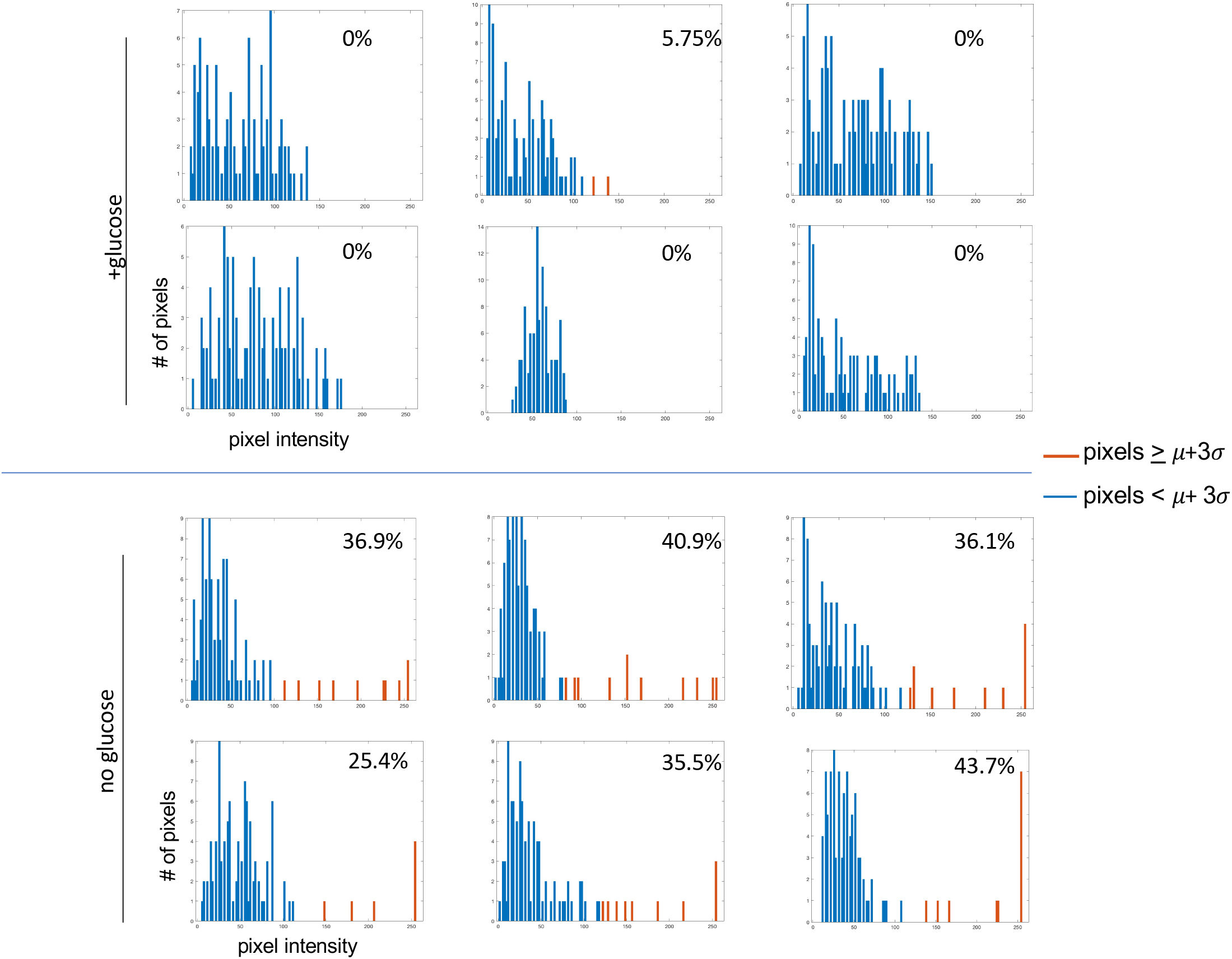
Scoring the distribution of GFP-Nab3 into granules in specific nuclei. Six nuclear masks consisting of 100 pixels were arbitrarily selected from images of strain DY4756 grown in the presence and absence of glucose. GFP-Nab3 pixel intensities for each mask were plotted in separate histograms. The pixels that exceed the μ+3σ threshold are highlighted in orange. The percentage shown in each histogram is calculated by:

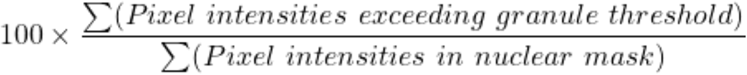

### Measurement of strain-to-strain differences in GFP-Nab3 granule-formation

Continuing to use our previous definition of a granule as a fluorescent spot with ≥4 contiguous 3σ pixels, we quantified the strain differences displayed in Fig. 1. The “high” granule forming strain observed above showed a penetrance of 86.3% (+/−5.7%) of the cells with a GFP-Nab3 granule (Fig. 6). The strain designated as “medium” possessed granules in 49.9% (+/− 13.7%) of its cells. The “low” strain had 3.5% (+/−0.1%) of its cells forming granules. As mentioned, the latter two were engineered identically to contain the two fluorescently-tagged proteins (Nab3 and histone H2B) in the lab strains BY4741 or W303, respectively. The highly efficient granule forming strain was derived from W303 and also contained the *NRD1* gene on a plasmid covering its deleted chromosomal copy. The granule forming efficiency of each strain was a reproducible property of individual clonal isolates of that strain and was maintained when cells were reseeded from thawed glycerol stocks. The molecular basis for this difference is thus far unclear, however the distinct phenotypes provide an avenue for a genetic analysis of granule formation.

**Figure 6.**
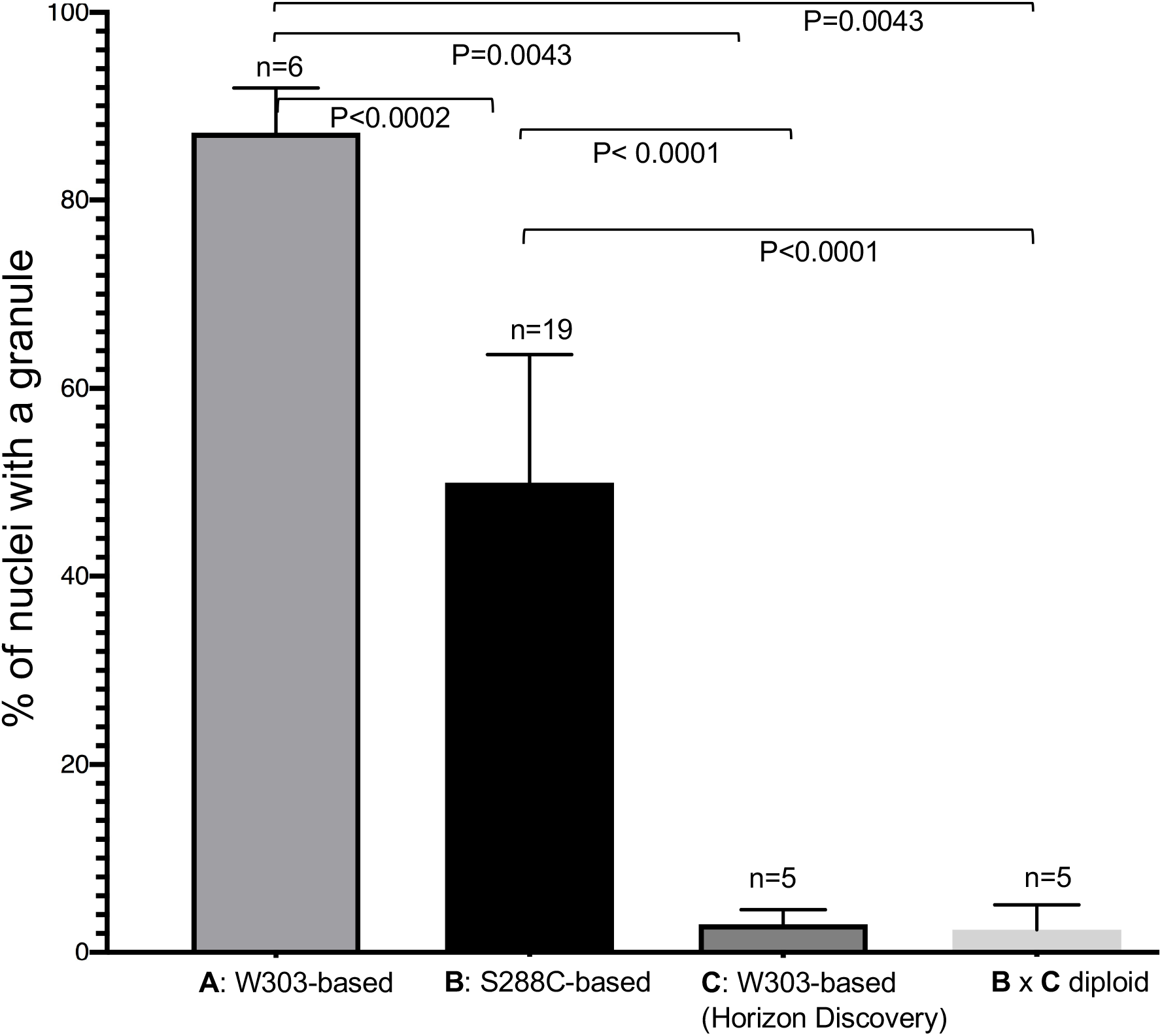
Different extents of granule formation in three distinct yeast strains The three yeast strains shown in Fig. 1 (DY4756, DY4580, DY4772) and the diploid strain resulting from the mating of DY4580 and DY47742 (DY4789) were glucose-starved and analyzed by fluorescent microscopy. Fields of cells from each were analyzed by the algorithm for biological replicates (n=6, n=19, n=5, and n=5 for strains DY4756, DY4580, DY4772, and DY4789, respectively) performed on different days. The average and standard deviation of the percent of cells containing GFP-Nab3 granules (≥ 4 adjacent 3σ pixels) were calculated for each strain and plotted on a bar graph. Fields contained an average of 1,210 cells each.

As an initial test of this idea, we mated the medium and low granule-forming strains to test which phenotype was dominant. Diploids were isolated and examined by confocal microscopy and the algorithm was applied to the digital images. Based on the number of pixels in the red fluorescent protein masks, the average size of the diploid’s nucleus was 1.7-fold larger than the average of the two haploids’, consistent with 1.7-fold larger total cell volume seen for diploids versus haploids (Mortimer, 1958). When granules were scored (Fig. 6), it was clear that the diploid showed the low efficiency version of the phenotype (2.4%+/−2.6%), suggesting that a low efficiency of granule formation is a dominant trait.

### The granule formation phenotype correlates with growth defects

The ability to identify strain differences in granule formation led us to consider other phenotypic differences that might lend insight into granule function, including growth temperature. Compared to the low efficiency granule forming strain, the high and medium granule formation phenotypes were associated with temperature sensitive growth at 37°C (Fig. 7A). The diploid that resulted from the mating of the medium and low efficiency granule-forming strains, grew as well at 37°C as its low efficiency granule-forming parent, consistent with the idea that the *ts* phenotype tracks with the low granule formation phenotype (Fig. 7A).

**Figure 7.**
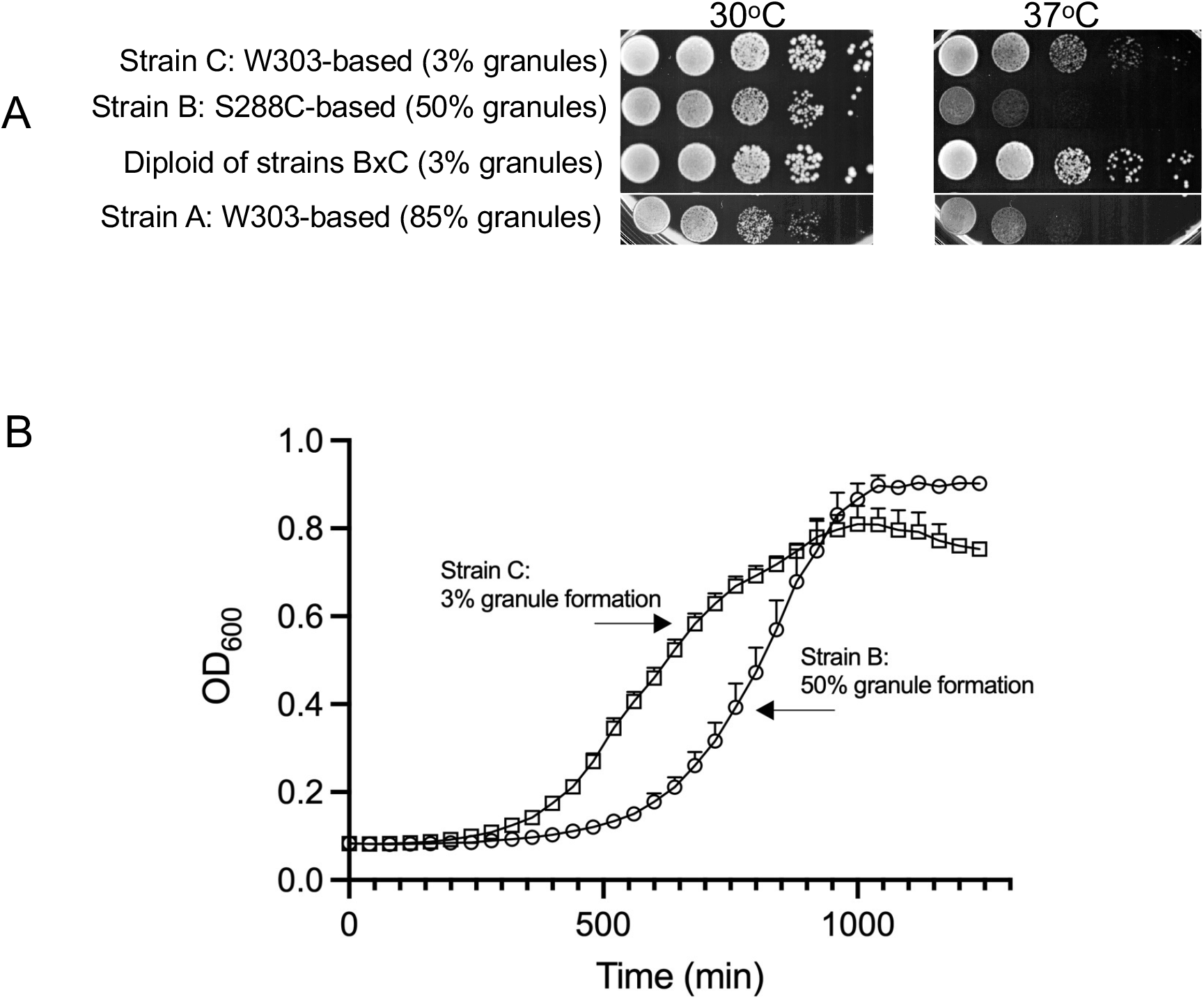
Growth defects are seen in high-efficiency granule forming strains. A. *Temperature sensitive growth*. Strains DY4756 (strain A), DY4580 (strain B), DY4772 (strain C), and the diploid strain DY4789 were diluted and spotted onto YPD and grown at the indicated temperatures. The medium and high-efficiency granule forming cells show reduced growth at 37°C. B. *Recovery from glucose starvation*. Strains DY4772 (strain C) and DY4580 (strain B) were washed free of glucose and incubated at 30°C for 2hrs before reseeding at OD_600_ = 0.1 in fresh glucose-containing medium. Cells were grown in 96-well plates at 30°C in a BioTek plate reader and light scattering (600 nm) recorded at intervals. Averages and standard deviations from biological triplicates were plotted. Strain B’s ability to resume growth in the presence of glucose is impaired relative to strain C’s.

When starved cells are refed, the granules rapidly resolve and cell division resumes (Loya et al., 2018). The different extents of granule formation between strains led us to consider if they might emerge from starvation at distinct rates. Cells were glucose starved and tested for how rapidly they restart growing after seeding them into fresh glucose-containing medium. The low efficiency granule-forming strain was relatively refractory to this stressor, attaining a mid-logarithmic density after 550 min in contrast to the medium efficiency granule-forming strain which took 800 min (Fig. 7B).

### The algorithm can measure filament formation by IMP dehydrogenase in cultured human cells

IMP dehydrogenase (IMPDH) is an enzyme involved in GTP metabolism that is highly conserved across species (Hedstrom, 2009). It detects substrate and product levels in cells and when nucleotide synthesis is needed, it possesses the ability to redistribute from its normal pan-cytoplasmic location to structures called rods and rings which contain organized polymers of the enzyme at its core (Carcamo et al., 2014; Schiavon et al., 2018). The consequence of this assembly is to render the enzyme more refractory to end product inhibition and to thereby increase total biosynthetic output of guanine nucleotides (Anthony et al., 2017; Calise et al., 2018; Duong-Ly et al., 2018). Most, if not all cells, can be induced to assemble IMPDH after treatment with the drug mycophenolic acid, which inhibits the enzyme, leaving cells depleted for guanine nucleotides (Carcamo et al., 2014).

To explore the versatility of the algorithm described above, we analyzed micrographs of human cells cultured from a T cell leukemia (Jurkat) treated with mycophenolic acid or vehicle and stained by indirect immunofluorescence with an anti-IMPDH2 polyclonal antibody (Fig. 8A). Nuclear DNA was counterstained with Hoechst 33342. Cells were examined with a BioTek Lionheart widefield microscope. Fluorescent images for each color channel were processed using the parameters described above with the Hoechst-fluorescence channel used to define a mask. Clusters of contiguous 3σ pixels in images labeled with anti-IMPDH2 fluorescence were scored in cells that were mycophenolic acid treated or untreated. As has been observed for a variety of cell types some rods pre-exist in the undrugged state and mycophenolic acid treatment strongly induces the extensive assembly of additional rods. (Carcamo et al., 2014).

**Figure 8.**
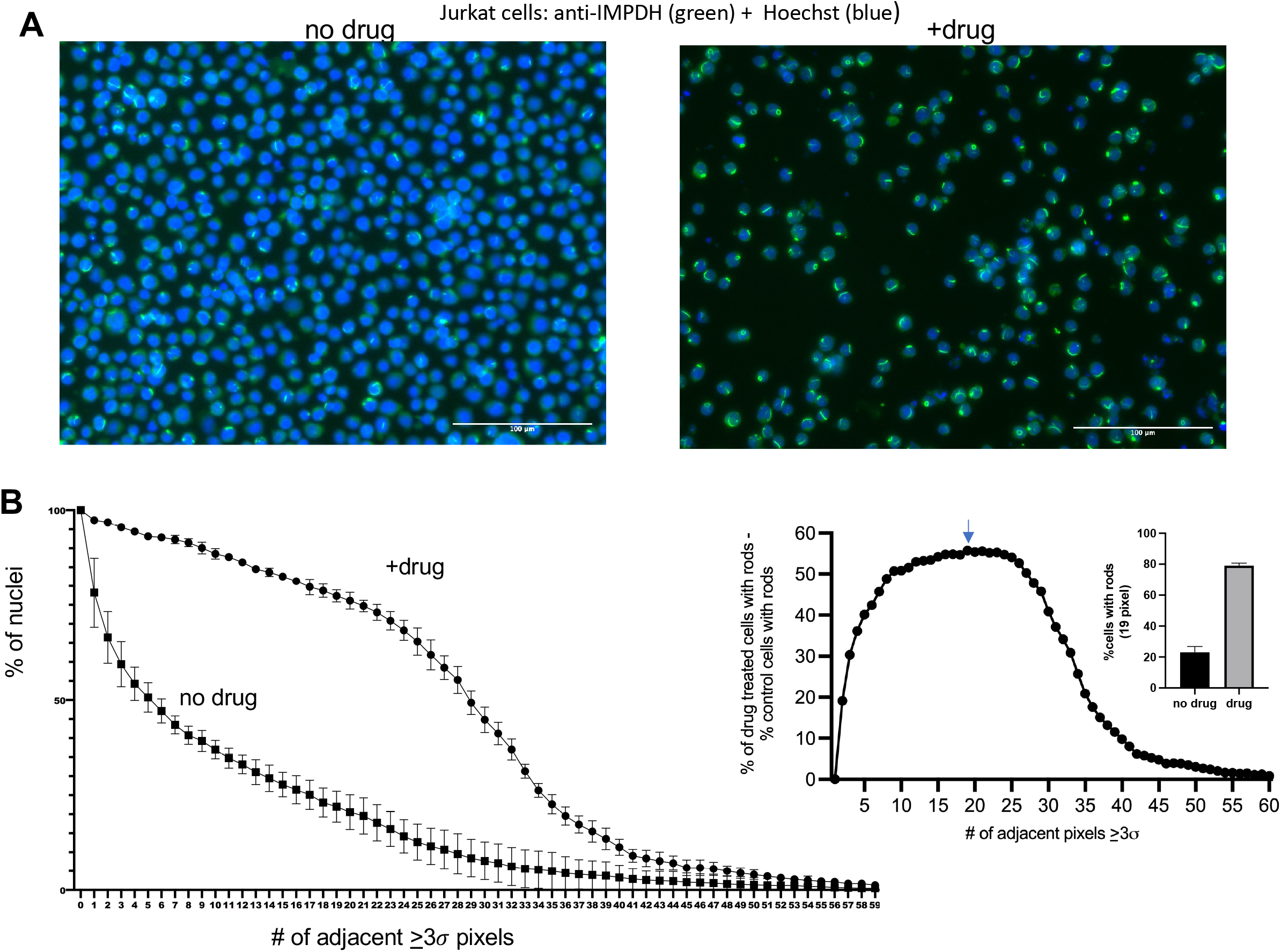
Quantification of drug-induced “rod and ring” formation by IMP dehydrogenase in human Jurkat cells. A. *Anti-IMPDH2 immunofluorescence*. Cultured Jurkat cells were treated with mycophenolic acid (right) or vehicle (left) and processed for immunofluorescence using an anti-IMPDH2 antibody. Nuclei were counterstained with Hoechst dye. Micrographs at 20x magnification are shown in panel A. B. *Quantification of drug-induced formation of IMPDH2 filaments*. Four fields (n=4) of Jurkat cells stained with anti-IMPDH2 were processed by the algorithm using Hoescht staining as a mask. The mean fraction of nuclei from these fields containing 0-59 adjacent 3σ pixels was calculated for drug- or vehicle-treated Jurkat cells, as indicated, and plotted (left panel). Error bars represent standard deviations. The difference between drug-treated and control cells was calculated for each number of contiguous 3σ pixels and plotted (center). The peak value (19 pixels) is indicated by the arrow. The average fraction of cells possessing 19 contiguous 3σ pixels (+/− standard deviation) was extracted from the graph on the left for drug-treated and control cells and is presented as a bar graph on the far right.

In examining a new protein in a different context, we again evaluated the appropriate size cutoff to score a granule. An application of the algorithm to these images showed that approximately half of the control cells display a condensate of IMPDH2 that is greater than or equal to five adjacent 3σ pixels (Fig. 8B). In contrast, just over half of the drug-treated cells showed 3σ pixel collections of 28 or more, consistent with the very large, micron-length filaments of IMPDH2 found in mycophenolate-treated cells (Carcamo et al., 2014). The clear difference between the size of GFP-Nab3 granules (above) and the large rods formed from IMPDH2, emphasize the algorithm’s versatility in its ability to set the extent of the area that defines a target compartment.

## DISCUSSION

We report the application of an algorithm used to show that distinct yeast strains display large differences in their elaboration of a nuclear granule composed of an essential RNA-binding protein. Characterization of a number of attributes of the granule became possible with this quantitative tool, including the fraction of the nuclear protein destined for the granule, the distribution of the number of granules per nucleus, the fraction of cells in a population that harbor a granule, the size of the granule, and the magnitude of the striking reorganization of the protein.

The scoring method is based on the fluorescent labeling of nuclei as a reference ‘mask’ and the identification of the changes in distribution of a protein of interest within that mask. Here we used the labeling of DNA or histones to provide the reference. Live cells with fluorescent proteins or fixed cells stained with antibodies, were equally good specimens. The process is largely automated, unbiased, and rapid and can yield data from large numbers of cells; in essence allowing rigorous determinations from ‘big data’. Using this tool, we could examine the extent to which a protein condenses from a pan-nuclear distribution into nuclear foci. We set a cutoff to define the size of a granule, calculate what fraction of the nuclear protein entered it, and show it was at the expense of the nucleoplasmic concentration of the protein. Generally, cells contained a single granule per nucleus; akin to a nucleolus and in contrast to the elaboration of a family of granules seen when stress granules are induced (Corbet and Parker, 2019). It was also possible to detect the difference in the size of nuclei between diploid and haploid cells by simply measuring the total number of pixels containing histone-coupled fluorescence.

Perhaps most importantly, we effectively quantified the fraction of cells in a population that contained a granule. With this we were able to identify clear differences between strains in terms of the penetrance of the granule-forming phenotype, and we showed that the low frequency of formation was dominant over a higher frequency in a diploid generated from both types of parents. This opens the way for future genetic analyses of the granule formation phenotype.

The algorithm also effectively identified the aggregation of IMPDH2 into the prominent rod structures seen in many cell types. Here a Hoescht-stained nuclear mask was used as a reference against which the relocation of the IMPDH2 enzyme was monitored by immunofluorescence. Although IMPDH2-based rods are most cytoplasmic, this was probably effective because the majority of the cellular volume of Jurkat cells is filled with the nucleus, so fluorescently stained IMPDH2 rods in the thin cytoplasmic region, which approximates a spherical shell, were largely super-imposed over Hoescht stained nuclei in two-dimensional maximum intensity projections. A future utility of the algorithm could be envisioned for other cell types with cytoplasmic compartments by employing a fluorescent pan-cytoplasmic reference protein against which a protein of interest can be studied.

In sum, we have developed and exploited an algorithm that can quantify the relative distribution and changes thereof, of proteins in fixed or living cells. The process is adaptable and available for community use through GitHub.

## MATERIALS AND METHODS

### Plasmid Construction

The plasmid pRS315NRD1 was generated by PCR amplification of Nrd1 from genomic DNA targeting the region 600 bp upstream and 500 bp downstream of the NRD1 open reading frame using oligonucleotides 5’-ATATAAGCTTCTACTGCGCACGTATAGCGC-3’ and 5’-ATATCTCGAGAATATGCCGGAAACGTCCAC-3’. The PCR product and pRS315 plasmid were cut with *Hin*dIII and *Xho*I restriction enzymes and ligated. The pRS315-2XHA-NRD1 plasmid was generated by inserting a 2XHA-tag after the start codon of the NRD1 ORF in pRS315NRD1 by PCR mutagenesis using the oligonucleotides 5’-TATCCGTATGATGTTCCGGATTATGCATATCCGTATGATGTTCCGGATTATGCACAGCAGGACGACGATTTTCA-3’ and 5’-CATTATGGGATGTTTAGTATGTTTGTTGC-3’.

### Yeast Strain Construction

Yeast strains used in this paper are presented in Table 1. DY4580 was generated using high efficiency lithium acetate transformation (Gietz et al., 1992) to transform DY4543 with a PCR product encoding *HTB2* tagged with mCherry and marked with *HIS3* that was amplified from genomic DNA of yeast strain YOL890 [gift from Dr. S Wente] using 5’-GCAGCTGAACCAGCTTTACC-3’ and 5’-GCTTTCAGTCGAAAACAGC–3’. Transformants were isolated by growth on SC his^-^ media and verified by PCR.

DY4724 was generated by transforming DY1620 with pRS315-HA-Nrd1. Transformants were selected on SC leu^-^ ura^-^ media. A single transformant was then struck onto 5-fluroorotic acid to cure DY4724 of pAC3314-Nrd1, resulting in DY4736. DY4736 was used to generate DY4746 by allele replacement using the pop-in/pop-out method of allele replacement (Rothstein, 1991) to integrate GFP onto *NAB3* as an N-terminal tag in its native chromosomal location. DY4756 resulted from transformation of DY4746 with a yomCherry-tagged *HTB2* PCR product generated from pFA6a-link-yomCherry-kan (Lee et al., 2013) using 5’-GGTGACGGTGCTGGTTTA-3’ and 5’-TCGATGAATTCGAGCTCG-3’.

DY4771 was generated using allele replacement using pop-in/pop-out to integrate the green fluorescent protein open reading frame in frame with *NAB3* in its native chromosomal location. DY4772 resulted from homologous recombination using a PCR product of mCherry-tagged *HTB2* into DY4771, as described above for the generation of DY4580.

The DY4789 diploid was generated by mating DY4580 and DY4772. Cells were mixed and selected on minimal media supplemented with uracil and leucine. The resulting colonies were then replica plated onto SC trp^-^ and SC met^-^ plates to select for potential diploids. Candidates were confirmed for diploid heterozygosity by PCR of *MET15* and *TRP1*.

### Yeast Growth Assays

Yeast strains were grown to saturation at 30°C in the appropriate selection media. For solid medium growth assays saturated cultures were diluted with sterile water to a concentration of 10^7^ cells/ml using 1 OD_600_ unit = x cells/ml, diluted serially 10-fold, and five spots of 10 μl were applied to the appropriate selective medium. Plates were grown at 24°C, 30°C and 37°C.

Cultures were inoculated and grown to saturation at 30°C. Optical densities (600 nm) were measured and cultures diluted to a final optical density of 0.1. Three 100µl biological replicates were plated in triplicate in a 96 well plate and placed onto a BioTek Epoch 2 plate reader at 30°C or 37°C. Optical densities were measured every 20 minutes until growth plateaued.

### Confocal Microscopy and Image Analysis

Cell containing GFP-tagged Nab3 and mCherry/yomCherry-tagged histone H2B, were grown to log phase in the appropriate, glucose-containing, selection media. Cells were split, pelleted, and washed three times into the appropriate media with or without glucose (starvation media). Cells were incubated at 30°C for 2 hours, pelleted and placed onto single cavity microscope slides containing 1.5% agar pads in the appropriate media. Z-stacks of both bright field and fluorescence were simultaneously obtained at room temperature using a Leica SP8 point scanning confocal microscope using the HyD (GaAsp) and PMT-transmitted light detectors, an HC PL APO 63X, 1.40 numerical aperture oil immersion lens, working distance 0.14mm. The microscope settings used are as follows:488nm line from an argon laser and 594 helium neon laser, pinhole 1 AU, image capture format 1024×1024, scan speed 400 Hz, pixel dwell time 1.2µs. Images were acquired using the Leica Application Suite X 3.0.2.16120 (Leica). Images presented in figures are displayed as maximum intensity projections. Color channels were split using FIJI (Schindelin et al., 2012) and quantified using the MATLAB algorithm (MATLAB., R2021a). The algorithm is available through GitHub.

*Jurkat cell culture*-Jurkat E6-1 cells were obtained from ATCC and grown in RPMI (Gibco) supplemented with 10% fetal bovine serum, penicillin and streptomycin in the presence of 5% CO2. Cells were maintained in T-75 flasks, passaged when ≤70% confluent, and routinely tested for mycoplasma contamination (Lonza). Cells were plated in 12-well plates at a concentration 10,000 cells/mL. Mycophenolic Acid (MPA; Sigma) dissolved in 0.1M NaOH was administered to cells at a final concentration of 10μM. Cells not destined for MPA treatment received an equal volume of the drug vehicle. Cells were treated for two hours, washed twice with phosphate buffered saline (PBS) supplemented with 10μM MPA (treated cells) or equal volume of vehicle (0.1M NaOH). Cells were resuspended in 167μL PBS with appropriate supplement and pipetted onto glass coverslips pre-treated with poly-L-lysine (Sigma; P4707) for 1 hr at 22 °C. Cells were then fixed with 4% (v/v) paraformaldehyde (Electron Microscopy Sciences; 15710) in PBS for 15 min followed by washing with PBS twice for 5 min. Cells were permeabilized with 0.1% Triton X-100 in PBS and incubated for one hour with 1% BSA (USBiological) in PBS. Cells were incubated overnight with rabbit anti-IMPDH2 antibody diluted 1:10,000. After 16 hrs, the coverslips were washed with PBS four times for 5 min each). The coverslips were then incubated for one hour with goat anti-rabbit AlexaFluor 488 (Invitrogen) antibody diluted 500-fold. The coverslips were washed with PBS twice for 5 min each followed by staining with Hoechst 33342 (Invitrogen), for four minutes. Coverslips were washed twice with PBS for 5 min each, and mounted on glass slides with a mounting mixture of 1, 4-phenylenediamine (Sigma) and Mowiol 4-88 (Sigma). The cured slides were imaged on a Biotek Lionheart FX using the “DAPI” and “GFP” filter cubes in addition to its brightfield capability. The three channels were merged in FIJI to generate a composite image.

## ACKNOWLEDGEMENTS

The authors acknowledge helpful discussions with, and assistance from, Drs. Judy Fridovich-Keil, Sohail Khoshnevis, and Homa Galei. The technical expertise of Laura Fox-Goharioon is also appreciated.

